# Sleep to remember, sleep to protect: increased sleep spindle and theta activity predict fewer intrusive memories after analogue trauma

**DOI:** 10.1101/2025.03.10.642388

**Authors:** Yasmine Azza, Mathias K. Kammerer, Hong-Viet Ngo, Mojgan Ehsanifard, Anna Wick, Klaus Junghanns, Ines Wilhelm-Groch

## Abstract

Recent evidence shows a strong correlative link between sleep disturbances and intrusive memories after traumatic events, presumably due to insufficient (nocturnal) memory integration. However, the underlying mechanisms of this link and the role of specific neural activities during sleep are poorly understood so far. Here, we investigated how the intra-individual affective response to an experimental trauma predicts changes in oscillatory activity during subsequent sleep and how these changes predict the processing of the experimental trauma. In a randomized within-subject comparison, twenty-two female, healthy participants (23.14 ± 2.46 years) watched a well-validated film clip including “traumatic” contents and a neutral film clip before bedtime on two separate nights. Heart rate was recorded during the film clips and nocturnal brain activity was recorded using 64-channel high-density EEG during subsequent nights. Intrusive memories were assessed via a seven-days diary and negative affect was assessed using laboratory trauma film reminders one week after the trauma film. An increased intra-individual heart rate during the trauma film predicted higher intra-individual sleep spindle amplitude the following night. Increased theta activity (4.25 - 8 Hz) during rapid eye movement (REM) sleep after the trauma film predicted fewer trauma film related intrusive memories and negative affect. Likewise, an increase in sleep spindles after the trauma film predicted fewer trauma film related intrusive memories. Our findings suggest that an experience-dependent up-regulation of these nocturnal oscillatory activity patterns, which are known to be involved in adaptive memory consolidation processes, serves as a protective factor against trauma-related intrusive memory development. Particularly, increased theta activity during REM sleep and sleep spindle activity seem to be of importance here.

## Introduction

Exposure to a stressful or traumatic event, such as interpersonal violence, accidents, or natural disasters, is a risk factor for many psychopathologies, with posttraumatic stress disorder (PTSD) being the most frequent mental disorder that may develop after trauma. Intrusive memories – the involuntary re-experiencing of a traumatic event often triggered by situational reminders – can emerge in the aftermath of a traumatic event. Recurring and distressing intrusions are not only a hallmark symptom of PTSD but are thought to be an early predictor for the development of PTSD or other psychopathological symptoms (Ehlers & Clark, 2000; Haag, Robinaugh, Ehlers, & Kleim, 2017). According to well-recognized etiology models of PTSD (dual representation theory (Brewin & Burgess, 2014); cognitive model of PTSD (Ehlers & Clark, 2000); emotional processing theory (Foa & Kozak, 1986)), intense arousal during the encoding of a traumatic event primarily stimulates perceptual and associative learning processes, which in turn can lead to an insufficient and fragmented memory trace of that experience. As a consequence, these unstructured memory representations may entail intrusive memories, which can be considered the result of a disruption in basic memory function (van Marle, 2015).

Given this, gaining deeper insights into mechanisms that drive these impaired memory functions after traumatic experiences seems key. One important factor in memory consolidation is sleep. Neural activity during sleep plays a crucial role in integrating new experiences into existing memory systems (Born & Wilhelm, 2012) and in reducing the intensity of distressing memories (Walker & van der Helm, 2009). Thus, it is not surprising that previous studies were able to show that sleep in the early aftermath of an analogue trauma can help reduce intrusive experiences and intrusion-related distress (Azza, Wilhelm, & Kleim, 2020; Kleim, Wysokowsky, Schmid, Seifritz, & Rasch, 2016; Porcheret et al., 2019; Wilhelm et al., 2021). Furthermore, decades of basic sleep science provide extensive evidence that slow-wave sleep activity, sleep spindle activity (brie bursts of neural activity in non-REM sleep (NREM)), and theta activity during rapid eye movement (REM) sleep are pivotal for successful memory consolidation. Additionally, most oscillatory activity during REM sleep can be attributed to the limbic system, a brain region known to be crucial for emotional memory consolidation (Genzel, Spoormaker, Konrad, & Dresler, 2015; Goldstein & Walker, 2014). However, there is growing evidence that neural activities during NREM and REM sleep complement each other in preserving and solidifying declarative aspects of an emotional memory, while at the same time attenuating its affective charge (Cairney, Durrant, Power, & Lewis, 2015; Rawson & Jackson, 2024). With regard to intrusive memories that emerge after traumatic experiences, these adaptive processes seem to fail. Thus, understanding the underlying mechanisms and roles of specific sleep oscillations that promote adaptive memory consolidation of traumatic experiences, and thereby potentially protect from intrusion development seems paramount in order to bring forth early and late interventions.

One way to investigate these underlying mechanisms is the use of an analogue trauma film paradigm. In this paradigm, highly distressing film content (e.g., interpersonal violence) is presented and elicited responses – which have been shown to be analogous to symptoms experienced after actual trauma (e.g., intrusive memories, physiological arousal, negative mood) – can be measured (Holmes & Bourne, 2008; James et al., 2016). Experimental sleep studies that used such trauma film paradigms have shown that increased slow-wave activity during a nap after film presentation was associated with less intrusion-related distress during the following week (Wilhelm et al., 2021). Furthermore, it has been shown that increased sleep spindle activity after film presentation was related to fewer intrusive memories (Kleim et al., 2016), to adaptive emotional memory processing (Kaestner, Wixted, & Mednick, 2013), and to improved sleep-dependent anxiety regulation (Natraj et al., 2023). Furthermore, a daytime nap *with* REM sleep (as opposed to without) after the exposure to an analogue trauma film entailed less subsequent intrusions (Wilhelm et al., 2021) and decreased the aversiveness of intrusive memories (Werner, Schabus, Blechert, & Wilhelm, 2021) in healthy individuals. Additionally, in a recent between-subject design study, greater theta activity during REM sleep after an experimental trauma was associated with significantly less intrusive memories (Sopp, Brueckner, Schäfer, Lass-Hennemann, & Michael, 2019). Further supporting the potential protective effects of theta activity, a cross-sectional study showed that individuals who experienced a traumatic event but did not develop PTSD, exhibited higher theta activity compared to individuals who developed PTSD (Cowdin, Kobayashi, & Mellman, 2014). However, these links need replication, have in parts not yet been studied in an overnight sleep study, and – most importantly – need to be investigated in a randomized within-subject design to inform about potential causality and inter-individual protective factors.

Given that only 10-20% of the people who experienced a trauma develop clinical post-traumatic symptomatology (Hidalgo & Davidson, 2000), this study aimed to determine potential protective factors against intrusive memory formation in a randomized within-subjects design in healthy individuals. For that, we used a validated, highly distressing film clip (i.e., “trauma” film) and a neutral film clip to examine how intra-individual changes in sleep physiology might protect from intrusion development. Firstly, we explored (i) the links between peri-trauma-film heart rate (as an indicator for arousal during encoding) and subsequent sleep oscillatory activity. Secondly, we expected (ii) increased EEG oscillatory activity in the slow wave, theta, and spindle spectrum after the trauma film exposure. Finally, we expected (iii) increased slow-wave activity, sleep spindle activity, and theta activity to be predictive of less trauma film related intrusive memories and negative affect.

## Methods

### Participants

Twenty-two healthy and female participants (mean age: 23.14 ± 2.46) took part in the present study and were recruited by the student mailing list of the University of Lübeck, Germany. Prior to the experiment the Ethics Commission of the University of Lübeck approved all study protocols and participants provided written informed consent. As a compensation, participants could choose to either receive a monetary incentive or course credits for their study program. Exclusion criteria were assessed in advance via a semi-structured telephone interview. These were: (1) experience of traumatic events involving interpersonal violence, (2) frequently watching of violent movies, (3) presence of a neurological or psychiatric disorder, (4) reported habitual consumption of alcohol or cannabis, (5) scheduled intake of medication influencing sleep during the study interval, (6) shift work. All participants were instructed to avoid caffeine on the day of the experiment.

### Procedure

In a randomized within-subject comparison, each participant spent three nights in the sleep laboratory including polysomnographic recordings. The first night (T0) served as an adaptation of sleep to the new environment (i.e., adaptation night). Before going to sleep, participants filled in informed consent and a baseline battery of questionnaires including demographic data, the WHO-Five Well-Being Index (World Health Organization, 1998), the Emotion Regulation Questionnaire (Gross & John, 2003), the Pittsburgh Sleep Quality Index (Buysse, Reynolds, Monk, Berman, & Kupfer, 1989), and the Edinburgh Handedness Inventory (Oldfield, 1971). On the second night (T1), according to a predefined randomization list, participants either watched a neutral or a highly distressing 12-minute film clip (see section “Trauma film paradigm”) in a darkened room using headphones. They were further informed that the film material they will see could contain violent and/or distressing scenes and that they are free to withdraw from the experiment at any point in time. The third night (T2) took place with a minimal time gap of seven days. It was conducted identically to T1 except that participants watched the film clip they had not been presented with before (neutral or trauma film). After the trauma film night, participants were instructed to document every intrusive memory that came to mind over the following seven days via an online questionnaire linked on their smartphone (i.e., intrusion diary). Additionally, participants were exposed to trauma film reminders seven days after the trauma film night in our laboratory and were asked to rate their subjective negative affect before and after the presentation of trauma film reminders (i.e., intrusion provocation task). All participants underwent both conditions on two separate nights (see also Figure 1).

**Figure 1.**
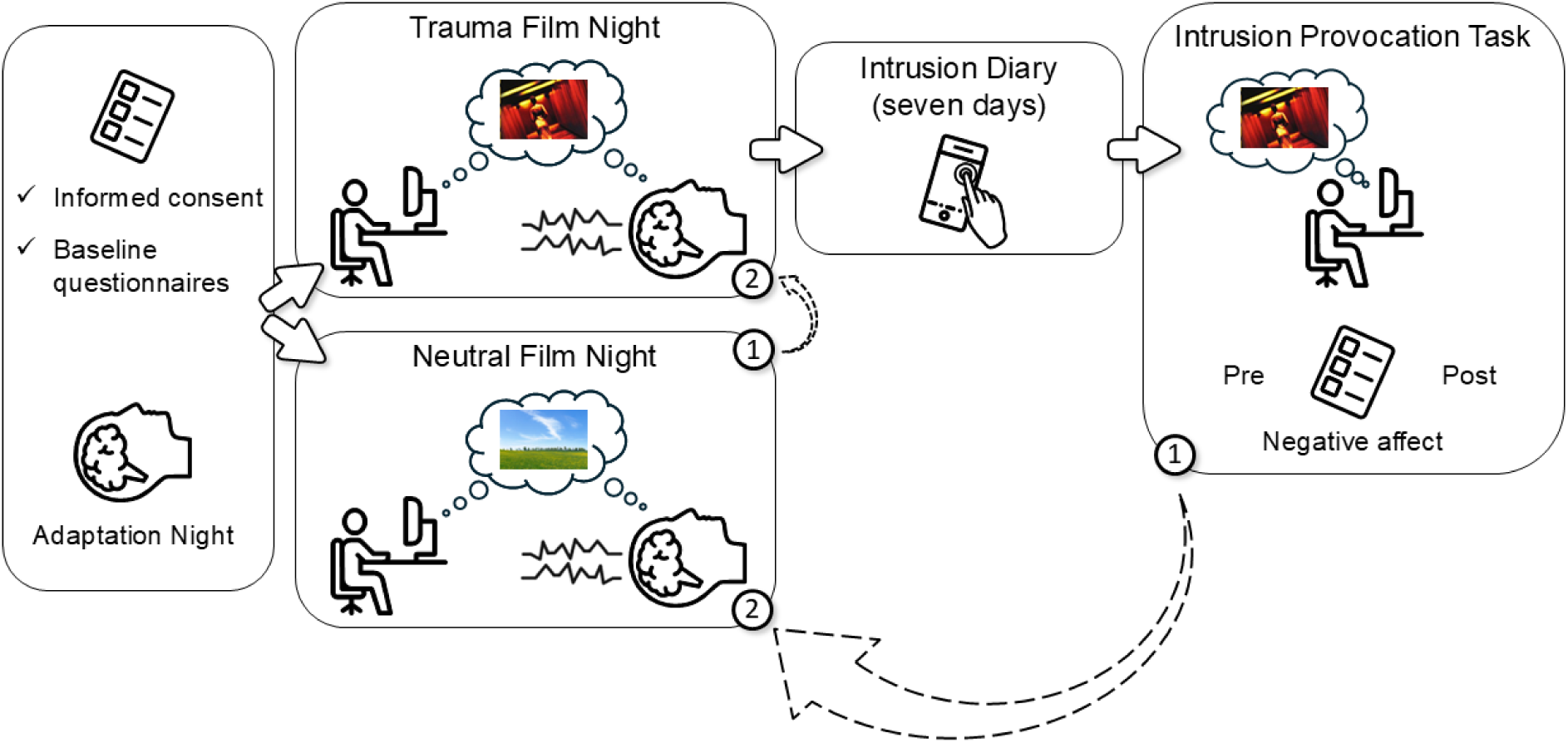
Study design. In a randomized within-subject comparison, participants watched either first a film clip including distressing contents (“Trauma Film”) or a neutral film clip (“Neutral Film”) before bedtime with polysomnography. All participants underwent both conditions on two separate nights (test-nights). For both film clips, heart rate and subjective arousal and mood were assessed. Intrusion diary and intrusion provocation task were only administered after trauma film night. There was a minimal time gap of two days between adaptation night and first test-night and a minimal time gap of seven days between each test-night. Numbers (1) and (2) indicate in what sequence an individual underwent the study procedure depending on randomized assignments.

### Trauma film paradigm^1^

The analogue traumatic stimulus in this study consisted of a previously used and validated 12-minute scene from the movie “Irreversible”, directed by Gaspar Noé including scenes of sexual violence (e.g., Kleim et al. (2016); Streb, Mecklinger, Anderson, Johanna, & Michael (2016)). Physiological and subjective reactions to the stimulus material were compared to a 12-minute neutral film depicting neutral social interactions as well as architectural scenes. Heart rate was recorded during both film presentations. Before and after both films, participants rated their mood and levels of arousal on a visual analogue scale, namely the Self-Assessment Manikins (SAM) (Bradley & Lang, 1994).

### Sleep EEG recordings and analyses

Brain activity during sleep was recorded using a Brain Vision system (LiveAmp) with a 64-channel electrode cap (BrainCap) and BrainAmp amplifiers (Brain Products, Munich, Germany). Two vertical electrooculogram (EOG) electrodes were placed above and below the left eye and two additional ones next to the lateral canthi to record horizontal eye movements. Three electromyography (EMG) electrodes (left and right upper chin and beneath chin) and two electrocardiography (ECG; left lower rib cage and right clavicle) electrodes were additionally applied. Impedances were kept below 10 kΩ. Brain activity was recorded with a sampling rate of 500 Hz.

For the analysis, the EEG recordings were re-referenced to mean activity of both mastoids and sleep stages were determined by visual scoring according to the American Academy of Sleep Medicine standard criteria (AASM; Iber, Ancoli-Israel, Chesson, & Quan (2007)) using the software SchlafAus 1.0 (developed by Steffen Gais, unpublished, University of Tuebingen, Germany). Data was further preprocessed by inspecting all channels of each person for possible distortions and marking them. Next, all data was high pass filtered at 0.1 Hz and beforehand defined bad channels were interpolated. Power spectra for slow-wave activity (SWA; 0.5 - 4Hz) were averaged over all of NREM2 and NREM3 episodes, whereas for theta activity (4.25 - 8Hz), spectra were averaged over all REM sleep intervals using Fast Fourier Transformation in Matlab 2016b (The MathWorks Inc., 2016). To estimate the individual increase of SWA after sleep onset, an SWA slope for each participant was calculated by subtracting the maximum two-minute-mean of SWA during the first sleep cycle from the initial two-minute-mean divided by the number of two-minute intervals. Discrete sleep spindle events (12 - 16Hz) were detected during artifact free intervals of NREM sleep 2 and 3 as described in Ngo, Fell, & Staresina (2020).

### Intrusion Diary

Intrusive memories were assessed via a mobile daily diary for seven days following the traumatic film clip via a questionnaire link installed on the participant’s smartphone. The time and date of the entries were recorded. Participants were asked to document the sudden occurrence of any intrusive memories i.e., images, thoughts or auditory stimuli related to the trauma film-clip in open text format. They were further asked to classify the type of memory (image, sound, thought) and to indicate the arousal and the degree of distress they experienced associated with the intrusion on a scale from 0=*not at all* to 100=*very distressing/aroused*. (adapted from Kleim, Graham, Bryant, & Ehlers (2013)). Only memories that occurred suddenly during the participants’ day were counted as intrusions. To ensure that, we implemented a control item that asked for the suddenness of each memory. Participants received daily email reminders in the evening to remind them of entering possible memories in case they forgot. The main outcome variable was defined as the absolute number of intrusive memories reported over the seven days after trauma film exposure.

### Intrusion provocation task

During an intrusion provocation task seven days after the trauma film presentation, five trauma-film-associated pictures (i.e., trauma film reminders) were presented for four seconds each (suit, dark street, stairs, underpass, and a pedestrian tunnel). After each of the pictures, during a one-minute interval, participants were asked to place their fingers on the spacebar, close their eyes, and press the spacebar for every upcoming intrusion. The total number of indicated intrusions for each image was calculated. Before and after the task, self-reported negative affect was evaluated using the Positive and Negative Affect Schedule - German Version (Krohne, Egloff, Kohlmann, & Tausch, 1996).

### Statistical Analysis

Descriptive statistics and comparisons of mean values were analyzed using R (R Core Team, 2017). To validate that the trauma film induced an increase in arousal and negative mood in our sample, paired t-tests (pre/post film presentation) for each of the conditions (trauma/neutral) were calculated. To examine potential differences of the topographical activity patterns between the two conditions more closely, cluster-based permutation testing was conducted for all electrodes and frequency bands of slow-wave activity (0.5 - 4 Hz), theta activity (4.25 - 8 Hz), and spindle activity (12 - 16 Hz). Cluster-based permutation tests were conducted using FieldTrip (paired samples t-test, 1,000 iterations; Oostenveld, Fries, Maris, & Schoffelen (2011)). The cluster test statistic was assessed against the Monte-Carlo permutation distribution to test the null hypothesis of no differences between the two conditions. Intra-individual change scores (trauma vs. neutral film) for heart rate and sleep measures were calculated and used for further analyses. We examined whether individual changes in heart rate during the trauma film (compared to the neutral film) predicted individual changes in the analyzed sleep measures using Spearman correlation analyses. To control for multiple comparisons, we applied cluster statistics based on the Monte Carlo method. Further, we examined whether individual changes in sleep measures after the trauma film (compared to the neutral film) predicted intrusive memories in the intrusion diary and negative affect in the intrusion provocation task using Spearman correlation (ρ). We set the threshold to control for family-wise error (FWE) to p = 0.05 (one-sided test).

## Results

### Effects of experimental trauma film on arousal, mood, and sleep architecture

In line with previous studies, the participants’ moods significantly deteriorated (*t* = 5.78, df = 21, *p* < .0001) while arousal levels significantly increased (*t* = −4.64, df = 21, *p* < .0001) by the presentation of the trauma film. For the neutral film, mood scores did not change significantly, whereas arousal levels significantly decreased (*t* = 2.74, df = 20, *p* = .031) from pre to post (see Table 1).

**Table 1.** Mood and arousal in response to the film material.

| | Pre Film<br>Mean ± SEM | Post Film<br>Mean ± SEM | $t(df)$ | $p$ |
| --- | --- | --- | --- | --- |
| Trauma Film |  |  |  |  |
| Mood (SAM) | **6.55±0.34** | **4±0.33** | **5.78(21)** | **<.0001** |
| Arousal (SAM) | **4±0.43** | **6±0.34** | **-4.64(21)** | **<.0001** |
| Neutral Film |  |  |  |  |
| Mood (SAM) | 6.9±0.32 | 6.86±0.28 | 0.22(20) | .825 |
| Arousal (SAM) | **3.57±0.51** | **2.57±0.40** | **2.74(20)** | **.013** |

SEM = standard error of the mean; SAM = Self-Assessment Manikin

We further examined whether there were any detectable differences in sleep architecture when comparing the trauma with the neutral film condition. Participants showed significantly increased sleep onset latencies (*t* = 2.73, df = 21, *p* = .013) after the trauma film while there was no difference between conditions in any other sleep variable (see Table 2). Interestingly, participants showed a flatter slope of SWA increase during the first sleep cycle across 41 channels primarily over the right hemisphere after the trauma film condition exposure (*t* = −3.54 – −2.08, df = 21, *p* ≤ .01).

**Table 2.** Sleep architecture after trauma and neutral condition.

| | Trauma film<br>Mean ± SEM | Neutral film<br>Mean ± SEM | $t(df)$ | $p$ |
| --- | --- | --- | --- | --- |
| Sleep onset latency (minutes) | **23.69±2.79** | **18.15±1.72** | **2.73(21)** | **.013** |
| NREM 1 (%)<sup>1</sup> | 4.3±0.55 | 3.93±0.38 | 0.84(21) | .412 |
| NREM 2 (%)<sup>1</sup> | 55.98±1.12 | 56.88±1.42 | -0.61(21) | .547 |
| NREM 3 (%)<sup>1</sup> | 13.37±0.8 | 13.85±0.96 | -0.58(21) | .570 |
| REM sleep (%)<sup>1</sup> | 21.24±0.87 | 20.89±0.89 | 0.56(21) | .579 |
| Total sleep time (minutes) | 474.59±8.35 | 473.98±9.13 | 0.89(21) | .891 |
| Wake after sleep onset (%) | 5.09±1.14 | 4.42±0.92 | 0.56(21) | .579 |

<sup>1</sup> Proportion of sleep stages relative to total sleep time; SEM = standard error of the mean; REM = rapid eye movement sleep; NREM = non-REM sleep stages.

### Effects of experimental trauma film exposure on neural sleep correlates

Average slow-wave activity (*t* = −2.15 – 1.87, *p* > .05) and sleep spindle activity (count: *t* = −1.45 – 1.42, *p* > .05; amplitude: *t* = −1.12 – 1.01, *p* > .05) during NREM sleep, as well as average theta activity (*t* = −1.41 – 1.99, *p* > .05) during REM sleep after trauma film exposure did not differ significantly compared to that after the neutral film (see Figures 2A - 2D).

**Figure 2.**
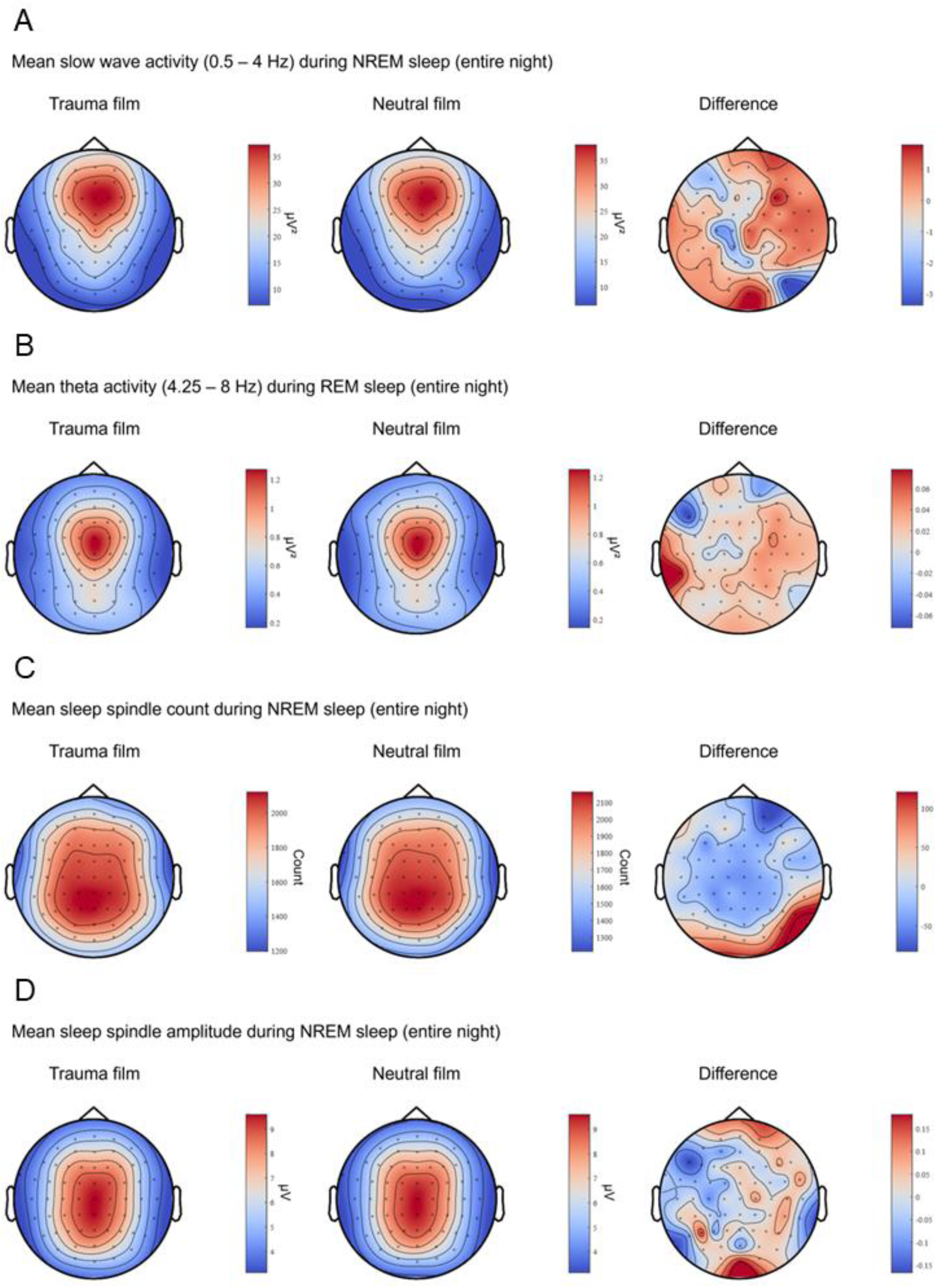
Differences in frequency bands comparing trauma and neutral film condition. Topographical distribution of the mean power spectra in three parameters of interest (slow-wave activity (A): 0.5 - 4Hz; theta activity (B): 4.25 - 8Hz; spindle count (C), and -amplitude (D): 12 - 16Hz) is illustrated for the trauma film (left), the neutral film condition (middle), and the difference between the two conditions (right). None of the measured frequency bands showed significantly different activities between trauma and neutral film conditions.

### Correlation between peri-trauma-film heart rate and sleep oscillations

Individual peri-trauma-film heart rate during the trauma film did not significantly predict slow-wave activity (see Figure 3A) or theta activity the following night (see Figure 3B). Intra-individually increased peri-trauma-film heart rate significantly predicted intra-individually increased sleep spindle amplitude the following night in 25 channels over frontal, central, parietal, and parieto-occipital areas (ρ = 0.586 – 0.404, *p* < .05; see also Figure 3D). No significant correlation was found for sleep spindle count (see Figure 3C).

**Figure 3.**
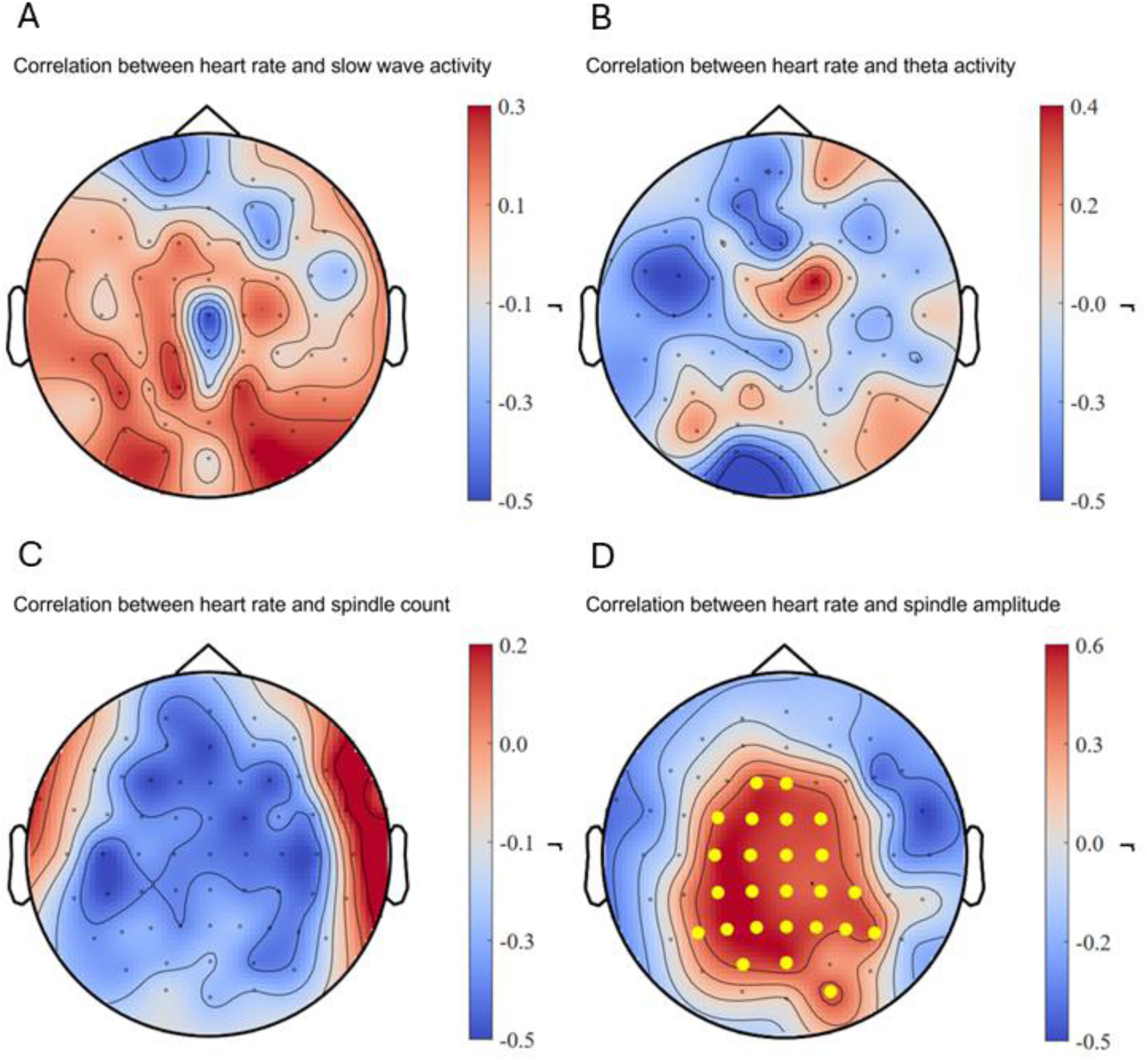
Correlation between heart rate during film exposure and sleep measures. (A-C) No significant correlations were found for slow-wave activity, theta activity, and sleep spindle count. (D) Intra-individual increase in heart rate during the trauma film (as compared to the neutral film) significantly predicted intra-individual increase in sleep spindle amplitude after the trauma film (compared to neutral). Dots in yellow mark the channels contributing to the significant cluster.

### Prediction of intrusions and affective response to trauma film reminders

Individual changes in slow-wave activity after the trauma film exposure did not predict intrusive memory formation during the seven days intrusion diary period nor did it predict negative affect during the intrusion provocation task (see Figures 4A and 5A). As hypothesized, an intra-individual increase in theta activity during REM sleep over left frontal, temporal, and parietal as well as right temporal and parietal areas in 30 channels after the trauma film significantly predicted less intrusive memories during the seven days intrusion diary period (ρ = −0.747 – −0.364, *p* < .05; see also Figure 4B). Similarly, intra-individually increased theta activity during REM sleep over frontal areas in 14 channels after the trauma film predicted less negative affect during the intrusion provocation task (ρ = −0.592 – −0.384, *p* < .05; see also Figure 5B). Intra-individually increased sleep spindle counts across nearly the entire scalp in 56 channels after the trauma film predicted significantly fewer intrusive memories during the seven days intrusion diary period (ρ = −0.672 – −0.366, *p* < .01; see also Figure 4C). Other individual changes in sleep spindle activity after the trauma film did not predict intrusive memories nor negative affect during the intrusion provocation task (see Figures 4D, 5C, and 5D).

**Figure 4.**
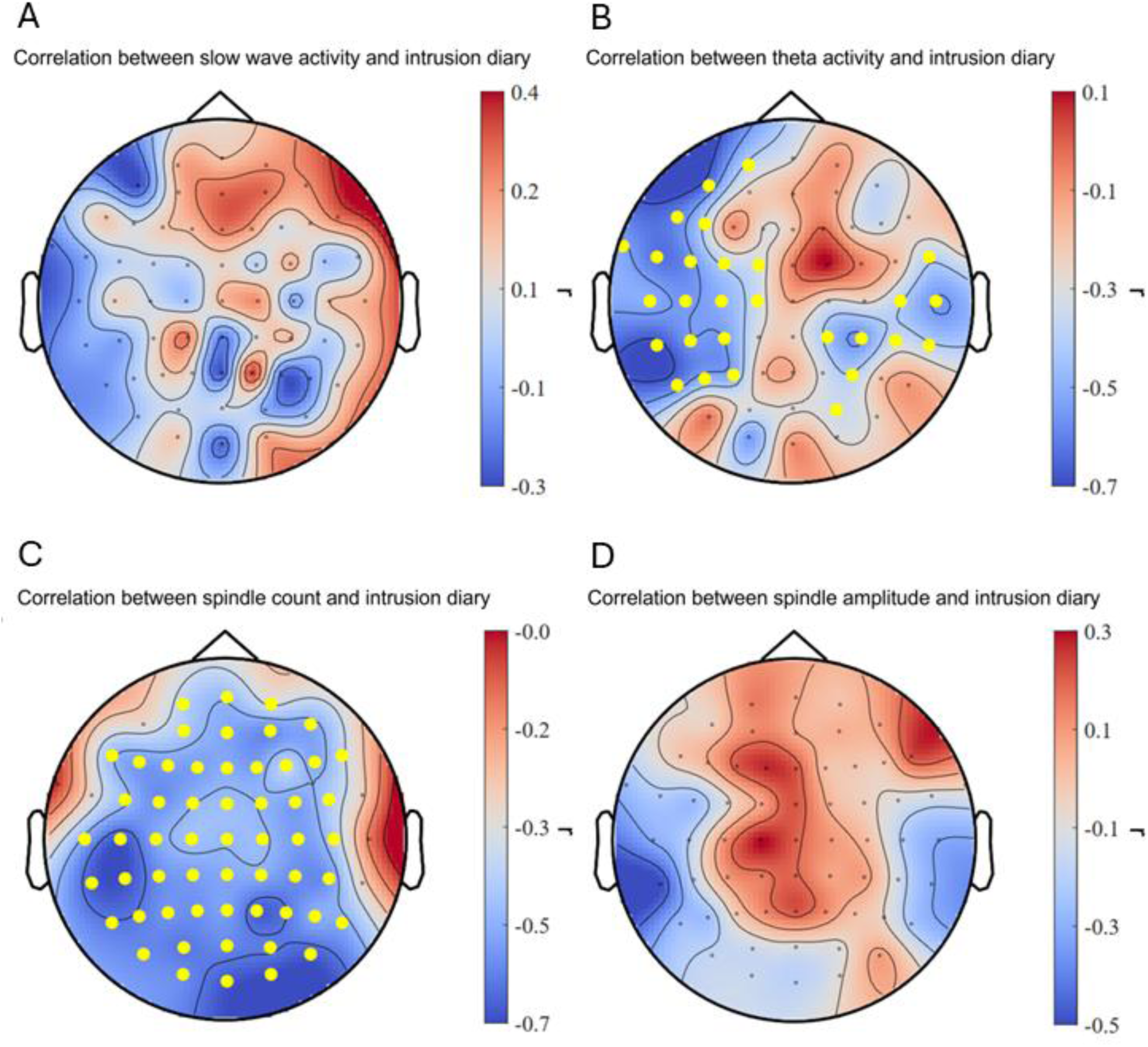
Correlations between intra-individual sleep changes and intrusions. An intra-individual increase of theta activity (B) and sleep spindle count (C) after the trauma film (compared to neutral) was significantly associated with less intrusive memories reported during the seven days intrusion diary period. No significant correlations were found for slow-wave activity and sleep spindle amplitude (A and D). Dots in yellow mark the channels contributing to the significant cluster.

**Figure 5.**
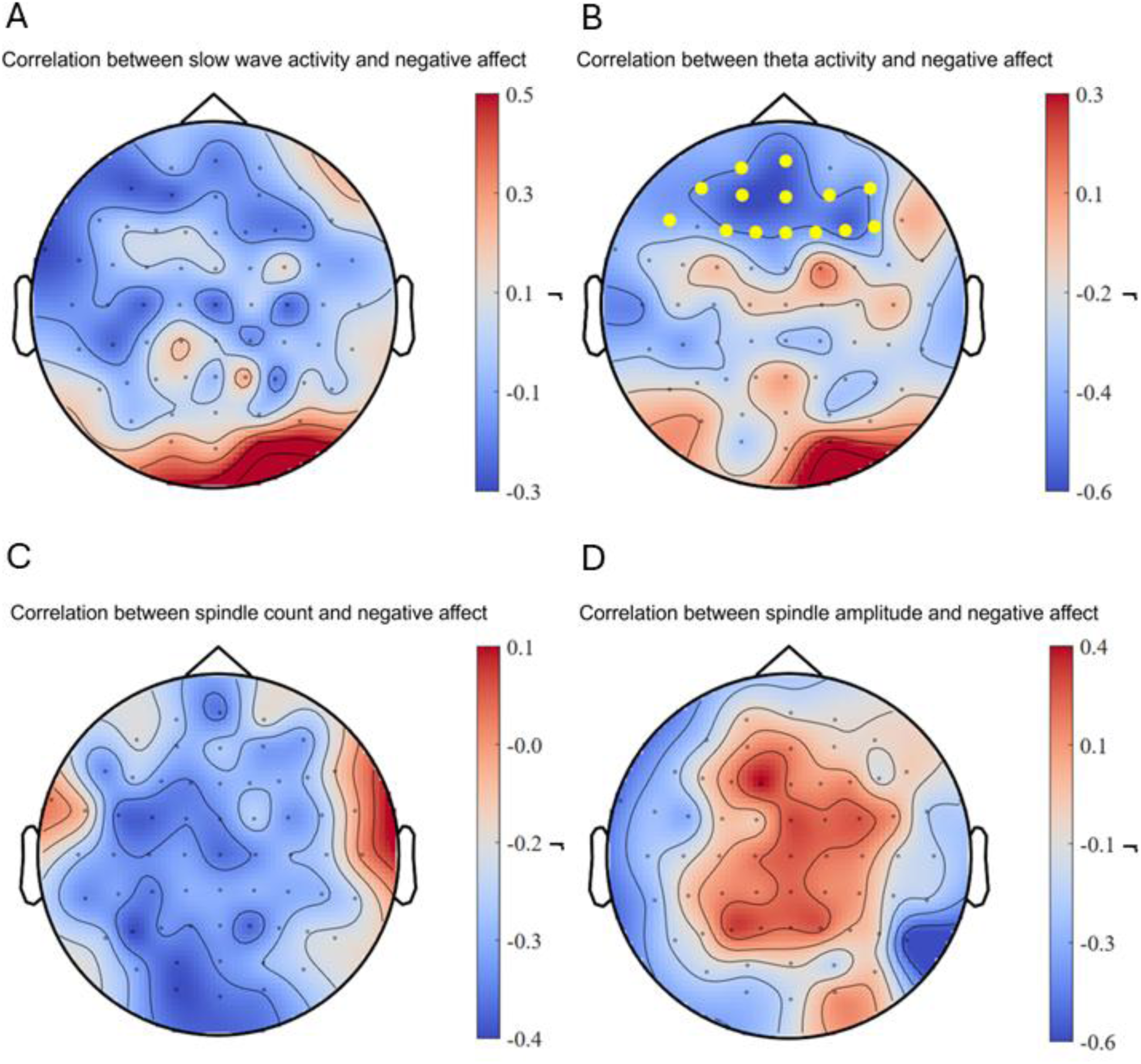
Correlations between intra-individual sleep changes and negative affect. An intra-individual increase of theta activity (B) after the trauma film (compared to neutral) was significantly associated with less negative affect during the intrusion provocation task one week after the trauma film exposure. No significant correlations were found for slow-wave activity, sleep spindle count and sleep spindle amplitude (A, C, and D). Dots in yellow mark the channels contributing to the significant cluster.

## Discussion

Here, we differentiated for the first time the impact of an experimental trauma film versus a neutral film on intra-individual neural sleep activity using high density EEG recordings in a within-subjects design. This allowed for conducting fine-grained cluster analyses of associations between intra-individual changes in sleep physiology and the processing of aversive experiences.

Against our hypothesis, we did not find significant changes in mean slow-wave activity, sleep spindle activity, and theta activity after the trauma film compared to the neutral film. This most likely can be attributed to high inter-individual differences on group level in response to a comparatively “mild” stressor. Since this study comprised only healthy participants, it is likely that they responded very differently to this “mild” stressor on an individual level. Consequently, the effects of the stressor may be present at the individual level but not at the group level.

However, in line with our expectations and with previous studies (Cowdin et al., 2014; Sopp, Brueckner, Schäfer, Lass-Hennemann, & Michael, 2019; Wilhelm et al., 2021), intra-individually increased theta activity during REM sleep after the trauma film predicted less intrusive memories in the intrusion diary and less negative affect in the intrusion provocation task. This strengthens the notion of theta activity being an affective “depotentiator” (Walker, 2009) and being actively involved in adaptive emotional memory consolidation (Nishida, Pearsall, Buckner, & Walker, 2009). Congruently, previous studies have shown that increased REM theta power significantly predicted enhanced emotional memory consolidation after a stress exposure (Kim et al., 2020). Thus, our findings support the notion that increased theta activity during REM sleep after a traumatic event is a protective factor against intrusive memories and trauma related negative affect. Moreover, our study is the first to demonstrate that theta activity is regulated in an experience-dependent manner and that this regulation is adaptive in face of a traumatic stressor.

Furthermore, in accordance with previous studies (Kleim et al., 2016; Wilhelm et al., 2021), we found an intra-individually increased sleep spindle count after the trauma film to predict less intrusive memories in the intrusion diary the following week. This aligns with research that showed increased sleep spindle activity is linked to reduced sleep-dependent anxiety responses in traumatized individuals (Natraj et al., 2023). Additionally, as sleep spindles change in response to experiences and promote synaptic plasticity (Fernandez & Lüthi, 2020), they play a key role in flexibly adapting to new demands. In sum, our findings suggest that increased sleep spindle activity facilitates adaptive emotional memory consolidation after traumatic events, helping to mitigate the development of intrusive traumatic memories (Natraj & Richards, 2023). However, individuals with PTSD often show increased sleep spindles, which has been suggested to reflect maladaptive over-consolidation of traumatic events and thus lead to more intrusive memories (van der Heijden et al., 2022). Considering these somewhat opposing findings, a differentiated assessment seems necessary. In light of the strong correlation between intra-individually increased heart rate during trauma film exposure and subsequent increased sleep spindle amplitude in this study, we propose temporarily increased sleep spindle activity after an acute stressor (accompanied by temporarily increased arousal) to reflect adaptive emotional memory processing. In contrast, increased sleep spindle activity in PTSD patients (accompanied by chronic hyperarousal) could reflect some sort of “dysfunctional replay” as a maladaptive attempt of the sleeping brain to integrate traumatic experiences into existing memory systems (see also Natraj & Richards (2023)).

Thus, one could speculate that moderately increased emotional reactivity (i.e., heart rate) during a stressful event promotes adaptive nocturnal emotional processing by up-regulating sleep spindle activity, which in turn would mitigate intrusion development. Corroborating this, previous studies have shown that reduced heart rate during a trauma film predicted the development of more frequent intrusive images in healthy individuals (e.g., Chou, La Marca, Steptoe, & Brewin (2014)) and that attenuated skin conductance responses (i.e., physiological arousal) during an emotion regulation task predicted later increased intrusion development in trauma-exposed individuals (Shepherd & Wild, 2014). This notion is further supported by a close affect-specific link between heart rate and neural activity in integral parts of the limbic system (Kuniecki, Barry, & Kaiser, 2003; Yang et al., 2007), a network crucial for memory consolidation (Catani, Dell’acqua, & Thiebaut de Schotten, 2013) and known to be altered in individuals with PTSD (Shin, Rauch, & Pitman, 2006). Generally speaking, when dealing with traumatic experiences, emotional engagement up to a certain degree as opposed to detachment (i.e., dissociation) has been shown to be beneficial in preventing or reducing PTSD symptoms, presumably by supporting adaptive (nocturnal) emotional memory processing (Möller, Söndergaard, & Helström, 2017; Rauch & Foa, 2006). However, it remains to be determined whether there are certain levels of arousal at different time points around a traumatic event that are beneficial for or detrimental to adaptive memory encoding and subsequent (nocturnal) consolidation (see e.g., Chou et al. (2014) for an attempt to decipher distinct effects of increased arousal at different time points around an analog traumatic event).

Strengths of our study include the conduct of a within-subjects design, which allowed us to assess and examine intra-individual changes in heart rate and sleep measures and thereby emulate real life inter-individual differences in response to traumatic events more accurately. Furthermore, instead of daytime naps, we used whole night high-density EEG recordings, which warrant higher spatial resolution and independence of ultradiane phases. As to limitations, our study comprised a relatively small sample with only female, non-clinical participants, which limits generalizability. Further, although analogue trauma film paradigms are widely used, they do not necessarily reflect real life trauma and our findings therefore ideally would need replication in naturalistic settings.

In conclusion, our findings provide further evidence for theta activity during REM sleep and sleep spindle activity during NREM sleep after analogue trauma to be up-regulated in an experience-dependent manner and by this being protective against intrusion development.

Furthermore, our findings suggest a close interplay between physiological reactivity during a traumatic event and subsequent sleep spindle activity. Given that sleep disturbances are prevalent in individuals with PTSD (Pace-Schott, Germain, & Milad, 2015) and are likely to contribute to PTSD symptom formation in the aftermath of traumatic events (Mellman, Pigeon, Nowell, & Nolan, 2007), interventions that aim to stabilize post-traumatic sleep quality seem paramount. Additionally, according to our findings, improving theta and sleep spindle activity after a traumatic event could promote adaptive emotional memory consolidation and thereby possibly prevent PTSD symptom formation (for theoretical reviews, see Murkar & De Koninck (2018); Mushtaq, Marshall, Ul Haq, & Martinetz (2024)).

## Acknowledgements

This work was supported by a grant of the Swiss National Science Foundation (SNSF, 10001C_179241). We thank Anna Wick, Fenja Rohrberg, and Lea Strelow for their ambitious help in data collection.

## Footnotes

1 We acknowledge that watching a movie scene with distressing contents does not necessarily reflect real life trauma.

